# Time series analysis of SARS-CoV-2 genomes and correlations among highly prevalent mutations

**DOI:** 10.1101/2022.04.05.487114

**Authors:** Neha Periwal, Shravan B. Rathod, Sankritya Sarma, Gundeep Singh, Avantika Jain, Ravi P. Barnwal, Kinsukh R. Srivastava, Baljeet Kaur, Pooja Arora, Vikas Sood

**Affiliations:** Department of Biochemistry, SCLS, Jamia Hamdard, New Delhi, India; Department of Chemistry, Smt. S. M. Panchal Science College, Talod, India; Department of Zoology, Hansraj College, University of Delhi, New Delhi, India; Humber College, Toronto, Canada; Delhi Institute of Pharmaceutical Sciences and Research, Delhi; Department of Biophysics, Panjab University, Chandigarh, India; Division of Medicinal and Process Chemistry, CDRI, Lucknow, India; Department of Computer Science, Hansraj College, University of Delhi, New Delhi, India

**Keywords:** SARS-CoV-2, Pearson Correlation, Hierarchical Clustering, Mutations, Protein Dynamics, Residue Correlation

## Abstract

The efforts of the scientific community to tame the recent SARS-CoV-2 pandemic seems to have been diluted by the emergence of new viral strains. Therefore, it becomes imperative to study and understand the effect of mutations on viral evolution, fitness and pathogenesis. In this regard, we performed a time-series analysis on 59541 SARS-CoV-2 genomic sequences from around the world. These 59541 genomes were grouped according to the months (January 2020-March 2021) based on the collection date. Meta-analysis of this data led us to identify highly significant mutations in viral genomes. Correlation and Hierarchical Clustering of the highly significant mutations led us to the identification of sixteen mutation pairs that were correlated with each other and were present in >30% of the genomes under study. Among these mutation pairs, some of the mutations have been shown to contribute towards the viral replication and fitness suggesting the possible role of other unexplored mutations in viral evolution and pathogenesis. Additionally, we employed various computational tools to investigate the effects of T85I, P323L, and Q57H mutations in Non-structural protein 2 (Nsp2), RNA-dependent RNA polymerase (RdRp) and Open reading frame 3a (ORF3a) respectively. Results show that T85I in Nsp2 and Q57H in ORF3a mutations are deleterious and destabilize the parent protein whereas P323L in RdRp is neutral and has a stabilizing effect. The normalized linear mutual information (nLMI) calculations revealed the significant residue correlation in Nsp2 and ORF3a in contrast to reduce correlation in RdRp protein.

## Introduction

The novel coronavirus first appeared in Wuhan, China in December 2019 and became a public health emergency of international concern. Since its emergence, the virus has caused catastrophe across the globe. This virus known as SARS-CoV-2 has infected nearly 486 million people and killed more than 6.1 million globally (*WHO Coronavirus (COVID-19) Dashboard*, n.d.). Out of the seven known coronaviruses, (HCoV-OC43, HCoV-229E, HCoV-HKU1, HCoV-NL63, SARS-CoV, MERS-CoV, and SARS-CoV-2) [1] SARS-CoV-2 is highly pathogenic to humans [2]. This virus has linear, positive-sense, single-strand RNA as genetic material which is about 30000 bp long and is encapsulated by the Nucleocapsid protein which is one of the four structural proteins other being Spike, Envelope, and Membrane proteins [3]. Once the virus gains entry inside the cell, two viral polyproteins namely ORF1a and ORF1ab are formed. These polyproteins are then cleaved by the viral proteases into sixteen non-structural proteins. These proteins initiate the process of viral replication and transcription. Apart from the viral non-structural proteins, SARS-CoV-2 encodes eleven accessory proteins that play a key role in viral pathogenesis [4].

Among the non-structural proteins of SARS-CoV, Nsp14 along with Nsp10 and Nsp12 plays a key role in maintaining the integrity of the viral RNA thereby resulting in a lesser number of mutations as compared to other RNA viruses [5,6]. Despite the fact that SARS-CoV-2 mutates at a slower pace, this virus has evolved into numerous variants since the onset of the pandemic in December 2019 [7]. The continuous evolution of SARS-CoV-2 has already dampened the efforts of the scientific community to design vaccines and antivirals against this virus [8]. Since mutations are one of the driving factors for virus evolution, various studies have been done to identify genomic variants among SARS-CoV-2. These studies have led to the discovery of a wide number of genetic variations, including missense, synonymous, insertion and deletion in the genomic sequences of SARS-CoV-2. The most common types of mutations along the viral genome were reported to be missense and synonymous in nature [9]. Although synonymous mutations may not have a direct impact on protein function, they do have consequences as they may alter codon usage, translation efficiency as well as binding kinetics of microRNAs. Furthermore, it was postulated that the mutations in the 5′UTR may alter the virus’s transcription and replication rates, as well as the folding of the genomic ssRNA sequences [10]. SARS-CoV-2 genome analysis have revealed a substantial mutation bias towards U which might be caused by improved immunogenicity, selection for greater expression, and better mRNA stability [11]. The viral transmission rates are rapidly increasing as it is evolving. For instance, a single mutation (D614G) in the Spike protein has been shown to increase the infectivity of SARS-CoV-2 [12]. As the viral genome is accumulating mutations, there can be a possible association among these mutations. It has been shown that co-mutations Y449S and N501Y in the Spike protein can lead to reduced infectivity and play a major role in disrupting antibody mediated virus neutralization [13]. This implies that mutations can have a synergistic effect resulting in enhanced viral fitness and immune escape. Therefore, immediate steps are required to identify the correlations and clusters among the mutations in the viral genome. Several studies have been published in this direction. A study conducted by Zuckerman et al. analysed 371 Israeli genomic sequences from February to April 2020 and identified correlations among mutations with that of known clade defining mutations [14]. Another study was conducted by Wang et al. where they analysed pairwise co-mutations of the top 11 missense mutations that were prevalent in the United States [15]. The 12754 SARS-CoV-2 genomes from the USA were analysed to obtain these 11 missense mutations.

In another study conducted by Rahman et al., the authors analysed 324 complete and near complete SARS-CoV-2 genomic sequences obtained from Bangladesh which were isolated between 30 March to 7 September 2020 [16]. They found 3037C>T was the most frequent mutation that occurred in 98% of isolates. This mutation is a synonymous one that were shown to co-occur with 3 other mutations including 241C>T, 144008C>T, and 23403A>G. In another study, Chen et al. analysed 261323 sequences of SARS-CoV-2 across the globe to identify the most common concurrent mutations among the top 17 mutations which occurred in more than 10% of the genomes under study [17]. The authors observed that there was a steady increase in the number of concurrent mutations as the COVID-19 pandemic progressed. The authors further showed that early M type genotype having two concurrent mutations evolved into WE1 with two additional concurrent mutations. WE1 further evolved into WE1.1 by incorporating three additional concurrent mutations.

However, some of these studies were performed with viral sequences obtained from a specific region, as well as focussed on the handful of top and missense mutations only. We hypothesized whether a similar trend could be observed with the genomic sequences of SARS-CoV-2 obtained from around the world. In order to gain a better understanding into the origin of mutations among SARS-CoV-2, we analysed viral genomic sequences in a time series-dependent manner. Meta-analysis of SARS-CoV-2 time-series data led us to the identification of highly significant mutations. We performed two highly common statistical tools including Pearson Correlation and Hierarchical Clustering to identify co-mutations occurring in viral genomes. In-silico protein dynamics were then used for the characterization of some of the highly prominent co-related mutations circulating in SARS-CoV-2.

## Material and Methods

### SARS-CoV-2 Genomic Sequences

Since the onset of the SARS-CoV-2 pandemic in 2019, the virus is continuously evolving resulting in the emergence of several variants. The availability of SARS-CoV-2 genomic sequences has been instrumental in understanding viral evolution and pathogenesis. To gain an in-depth understanding of the mutational landscape of SARS-CoV-2, we sought to analyse SARS-CoV-2 genomic data in a time-series manner. All the SARS-CoV-2 genomic sequences were collected in a month-wise manner (based on the sample collection month) from the Virus Pathogen Resource (ViPR) database [18]. In the current study, SARS-CoV-2 sequences were collected from January 2020 to March 2021 totalling to around 59541 samples.

### Meta-analysis of SARS-CoV-2 genomic sequences

Once the SARS-CoV-2 sequences were obtained and categorized by the collection month, we performed the meta-analysis of these sequences to identify highly significant mutations among circulating genomes. The genomes from various months were analysed with respect to the genomes collected in January 2020. The genomes obtained during the initial phase of infections tend to be close to the wild type with a few mutations as compared to the genomes collected at the later stages of the infection. For the identification of highly significant mutations, Meta-data-driven comparative analysis tool for sequences (META-CATS) was used [19]. All the analyses were performed with default settings and mutations having p>0.05 were considered significant. We obtained highly significant mutations for each month. Notably, SARS-CoV-2 genomic sequences collected in December 2020 did not yield any significant mutation. Therefore, December 2020 genomes were not included for further analysis.

### Correlation Coefficient

#### Pearson Correlation coefficient

Our analysis identified highly significant mutations some of which were common among several months. In order to identify the correlation among highly significant mutations in SARS-CoV-2 circulating genomes, we used the Pearson Correlation. We created an empty matrix with 30000 columns and 55759 rows using the NumPy module of Python. In this matrix, the number of columns represents the maximum possible length of the SARS-CoV-2 genomic sequences that were used in the study, while the number of rows represents the number of SARS-CoV-2 genomic sequences that we analysed. Once the empty matrix was created, for each genome of a particular month, we inserted the number “1” at those positions where Meta-CATS identified a highly significant mutation for that month and “0” was inserted at those positions where no mutation was identified. This task was performed using an in-house Python script. Therefore, we obtained a binary 55759×30000 matrix where the number “1” represented a position where a significant mutation was identified whereas a number “0” represented a position where no mutation was present. Once this matrix was prepared, we applied Pearson Correlation on this matrix to find out correlation coefficient among the highly significant mutations. We used the corr method of the pandas library to implement Pearson correlation.

### Clustering

#### Hierarchical Clustering

To further validate our results obtained from the Pearson correlation, we used another statistical tool called Hierarchical Clustering which groups the same objects in clusters thereby pointing towards the closeness of the objects. Hierarchical Clustering is a very computationally intensive process. Since highly frequent mutations tend to play a critical role in the evolution of the virus, we used the top 25 mutations that were present in >10% of the genomes used in this study [17]. Hierarchical Clustering was performed on the binary matrix of the top 25 mutations. We applied the figure_factory (create dendrogram) method of plotly that performs Hierarchical Clustering on the matrix and created a dendrogram that represents the highly significant mutations which are clustered together. To make the visualization more effective, we represent the dendrogram with a heatmap using the pdist and squareform method of scipy library. Values on the colour bar of dendrogram with heatmap correspond to distances between highly significant mutations. All these analyses were performed in python.

### Protein structure modelling and preparation

Our analysis identified nine highly significant and frequent mutations that were correlated with each other. We then sought to use computational tools to identify the effect of these mutations on protein structures. Out of these nine mutations, one mutation (241C>T) was present in 5’UTR which does not get translated into the protein. Two mutations (3037C>T and 28882G>A) are synonymous in nature and hence were not included for further analysis. Out of the remaining six mutations, we performed the detailed analysis of three mutations (23403A>G, 28881G>A, and 28883G>C) in our recent study [20]. Therefore, in this study, we targeted the remaining three mutations 1059C>T (T85I), 14408C>T (P323L) and 25563G>T (Q57H) to probe the impact of these mutations in Non-structural protein 2 (Nsp2), RNA-dependent RNA polymerase (RdRp) and Open reading frame 3a (ORF3a) respectively. The crystal structures of these proteins are available on RCSB Protein data bank (PDB) (https://www.rcsb.org/) but they have missing residues. Hence, we employed a deep learning-based protein modelling tool, RoseTTAFold [21] available at https://robetta.bakerlab.org/ to add missing residues in our proteins. Nsp2 (PDB: 7MSW) is 638 amino acids long and in crystal structure, the first three residues at the N-terminus are missing. RdRp (PDB: 7CYQ) is 942 amino acids long and 1-3 amino acids at N-terminus and 930-942 amino acids at C-terminus were missing. ORF3a (PDB: 6XDC) is only 284 amino acids long but a large number of residues from both the terminals (1-39 a.a. at N-terminal & 239-284 a.a. at C-terminal) and six residues (175-180 a.a.) were missing inthe protein structure. The RoseTTAFold modelled all these missing residues in three proteins except 9 histidine residues at the C-terminal in RdRp. These three modelled proteins were further analysed for mutation effects.

### Functional impacts of mutations

To investigate the effect of mutation on protein function, we used the widely popular PredictSNP web server [22] available at https://loschmidt.chemi.muni.cz/predictsnp/. This web tool is composed of six different predictors, PhD-SNP, MAPP, SNAP, PolyPhen-1, SIFT and PolyPhen-2 to predict whether mutation is deleterious or neutral. PredictSNP gives a consensus prediction score using these six predictors. These six predictors make use of different methodologies to predict the nature of mutation. PhD-SNP, MAPP, SNAP, PolyPhen-1, SIFT and PolyPhen-2 apply support vector machine, physicochemical characteristics and protein sequence alignment score, neural network approach, expert set of empirical rules, protein sequence alignment score and naïve Bayes respectively [22]. To calculate PredictSNP score, following equation is employed,

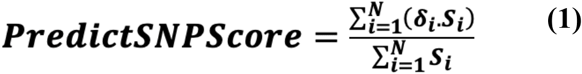

Where, *δi* is an inclusive prediction (neutral: -1 & deleterious: +1), *S*_*i*_ indicates the transformed confidence scores and N is the number of predictors. PredictSNP consensus score is between - 1 and +1, where -1 to 0 is designated for neutral and 0 to +1 for deleterious mutation.

### Effects of mutation on protein dynamics

The Normal mode analysis (NMA) based DynaMut [23] web tool (http://biosig.unimelb.edu.au/dynamut/) was utilized to probe the effects of a single mutation in each protein on protein stability and flexibility. The folding free energy change (ΔΔG) was calculated to exactly predict the stability of protein under the influence of mutants. In addition to its own ΔΔG prediction, DynaMut also predicts ΔΔG using NMA based ENCoM (Elastic network contact model) [24] and, other structure-based predictors like mCSM [25], SDM [26] and DUET [27]. The free energy change measures the energy difference between WT and MT proteins and gives insight to the stability of proteins. Additionally, DynaMut employs ENCoM to predict vibrational entropy energy (ΔΔS_vib_). The values of ΔΔS_vib_ are calculated for WT and MT by screening their all-atom pair interactions.

We utilized protein sequence-based Single amino acid folding free energy changes-sequence (SAAFEC-SEQ) [28] to validate the DynaMut predictions for WT and MT proteins. This tool is available at http://compbio.clemson.edu/SAAFEC-SEQ/ and uses different protocols such as protein sequence properties, evolutionary details and physicochemical properties to calculate ΔΔG value.

### Linear mutual information (LMI)

To understand the dynamical nature and fluctuations of biomolecules, Dynamical cross-correlation (DCC) and Linear mutual information (LMI) are employed widely [29–32]. Since DCC cannot calculate the correlation of concurrently moving atoms in perpendicular directions [33], we applied normalized LMI (nLMI) to overcome this limitation. To calculate the nLMI of WT and MT proteins, we employed Python-based correlationplus 0.2.1 tool [33]. To calculate nLMI, we used pdb files as input to the program. During the calculation, the program used the Anisotropic network model (ANM) to generate 100 models of WT and MT proteins and correlation was obtained using these models. Also, we calculated the difference in correlation between WT and MT proteins. To calculate nLMI between residue *i* and *j*, following equation is used;

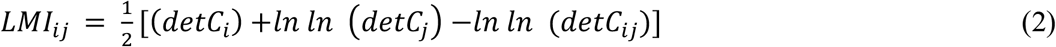

Where, 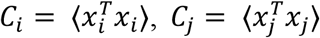, and *C*_*ij*_ = ⟨(*x*_*i*_,*x*_*j*_)^*T*^(*x*_*i*_,*x*_*j*_)⟩. And, *x*_*i*_ = *R*_*i*_ − ⟨*R*_*i*_ ⟩ and *x*_*j*_ = *R*_*j*_ − ⟨*R*_*j*_ ⟩ where, *R*_*i*_ and *R*_*j*_ are the atom *i* and *j* position vectors. In nLMI calculation, the LMI was considered greater than or equal to 0.3 and distance threshold was less than or equal to 7 Å. The 0 and 1 value indicate no correlation and complete correlation of residues respectively.

## Results and Discussion

### SARS-CoV-2 genomes

Since the onset of the COVID-19 pandemic, several studies have identified mutations in SARS-CoV-2 circulating genomes. However, in order to identify highly significant mutations, their origin and frequency, we analysed SARS-CoV-2 genomic sequences in a time-series manner. Since there can be a considerable time lapse between sample collection and sample processing, therefore, we used the sample collection date criteria to classify the SARS-CoV-2 genomic sequences to a particular month. We included a total of 59541 SARS-CoV-2 genomes that were collected from January 2020 till March 2021 (Table 1). Global distribution of the samples revealed that the majority of the samples were from the USA followed by Australia and India (Figure 1).

**Table 1:**
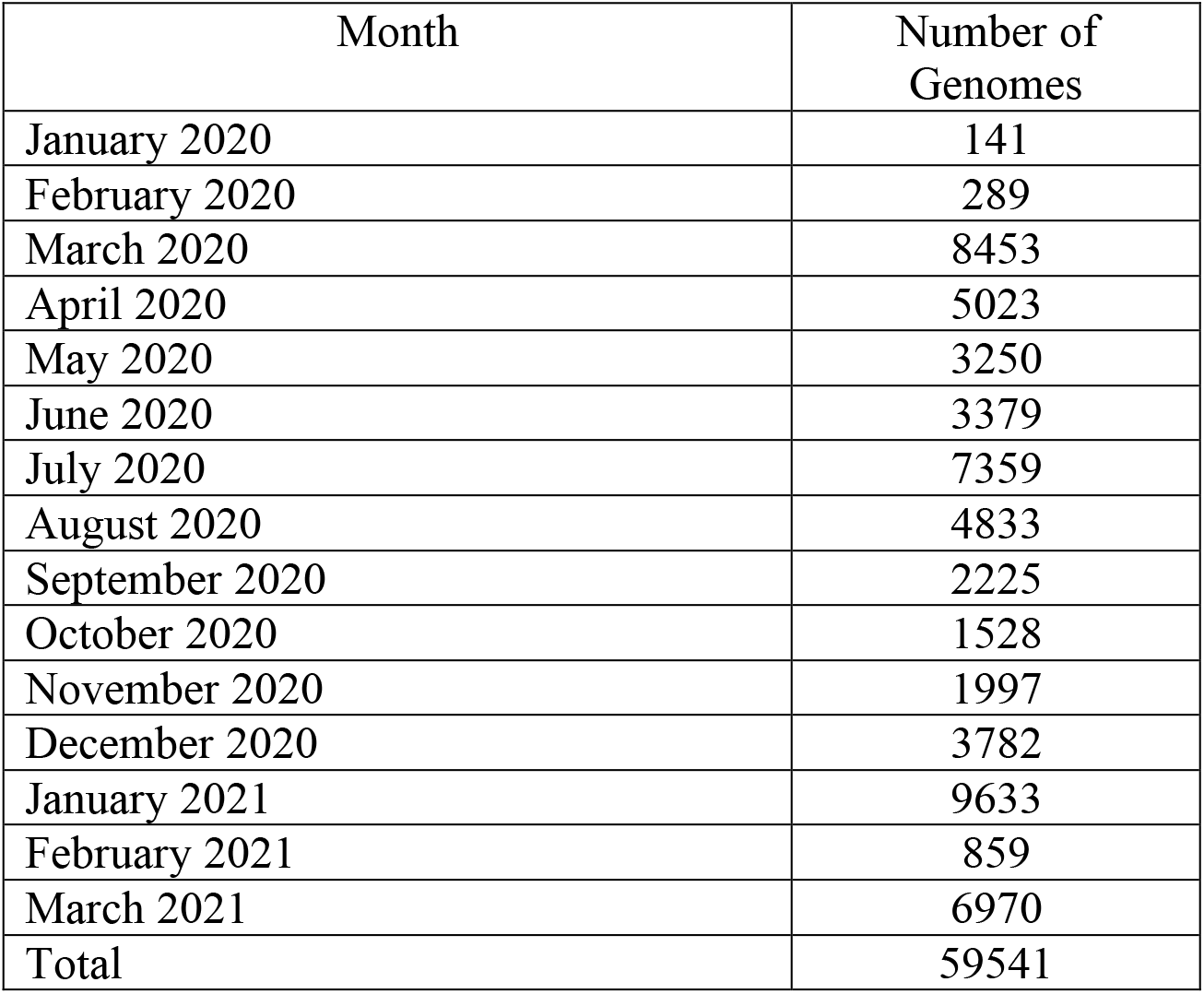
Table showing the number of genomes that were analysed in this study. The number of genomes collected in month wise manner is also shown in the table.

| Month | Number of Genomes |
| --- | --- |
| January 2020 | 141 |
| February 2020 | 289 |
| March 2020 | 8453 |
| April 2020 | 5023 |
| May 2020 | 3250 |
| June 2020 | 3379 |
| July 2020 | 7359 |
| August 2020 | 4833 |
| September 2020 | 2225 |
| October 2020 | 1528 |
| November 2020 | 1997 |
| December 2020 | 3782 |
| January 2021 | 9633 |
| February 2021 | 859 |
| March 2021 | 6970 |
| Total | 59541 |

**Figure 1:**
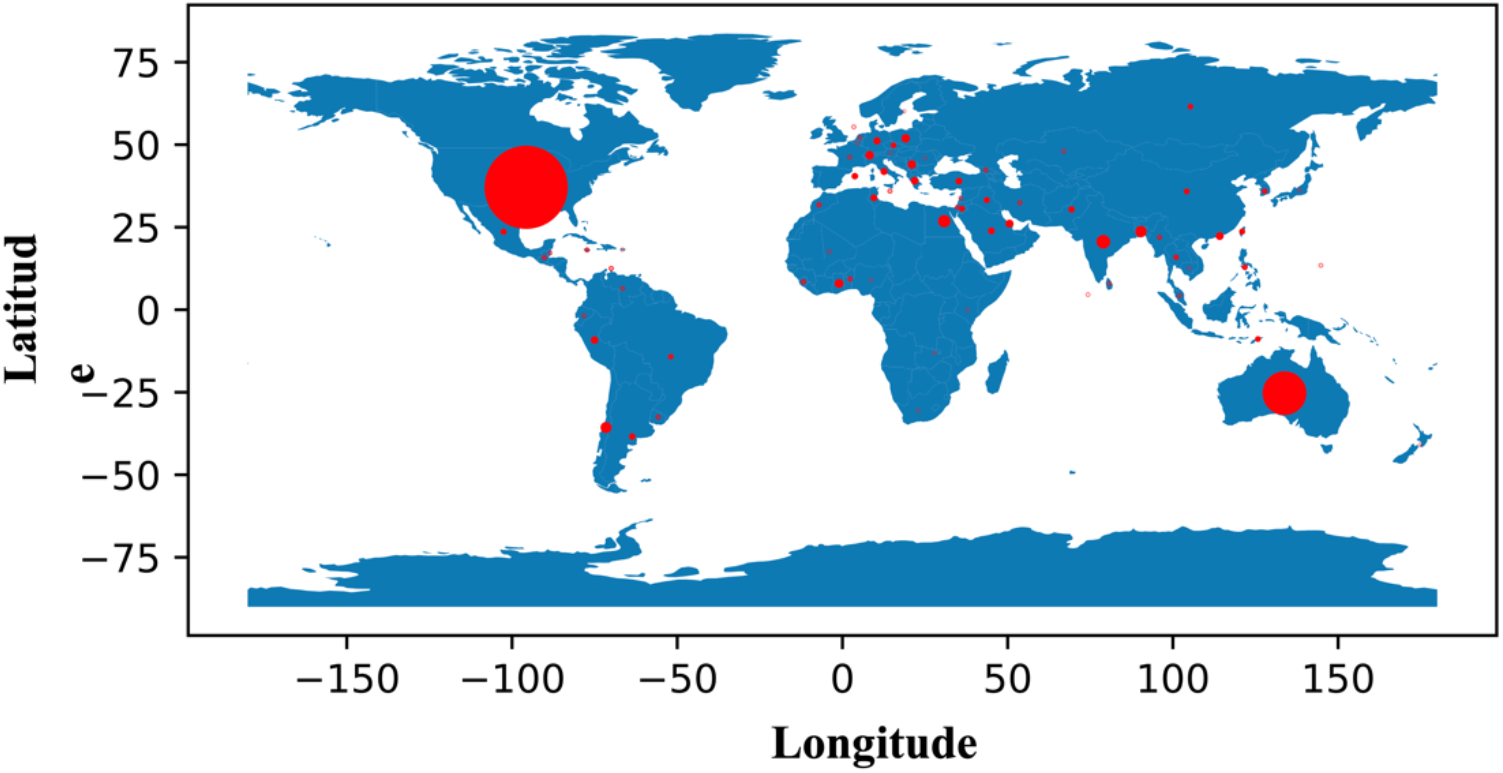
representing the geographical distribution of samples used in this study. Circle size indicate then number of samples from each geographical location. X-axis represents the longitude whereas Y-axis represents the earths latitude.

### Identification of highly significant mutations in SARS-CoV-2 genomes

Once the SARS-CoV-2 genomic sequences were grouped on the basis of the month, we used the META-CATS algorithm to identify significant mutations among the genomes. This algorithm compares two different datasets to identify highly significant mutations among them. As the genomic sequences collected at the start of the pandemic tend to be very similar to the parent sequence, hence all the SARS-CoV-2 genomic sequences collected in the month of January 2020 were grouped as control sequences. These control sequences were then analysed against the sequences obtained in the subsequent months to identify highly significant mutations in SARS-CoV-2 genomes of that month. We obtained highly significant mutations for each month except December 2020 (Figure 2(A)). Since mutations at the nucleotide levels might not lead to the change in amino acids, therefore we focussed our attention on the mutations at the amino acid level Figure 2(B-I). Across all the months we identified 940 unique mutations at the level of amino acids which were unevenly distributed among the genomes of SARS-CoV-2. Our analysis identified 610, 256, 33, 2, 11, 10, 16 and 2 mutations in SARS-CoV-2 ORF1ab, S, ORF3a, M, ORF6, ORF8, N and ORF10 proteins respectively. Since the length of SARS-CoV-2 proteins is highly variable, therefore we calculated the frequency of highly significant mutations at amino acid level in order to understand its distribution in the viral proteins. We observed that Spike protein had the highest frequency of mutations (20.10) followed by ORF6 (18.0) and ORF1ab (8.59) (Table 2). We observed that the membrane protein of SARS-CoV-2 had the least number of mutations as compared to the other proteins suggesting that this region among SARS-CoV-2 was highly conserved. The emergence of SARS-CoV-2 mutant named Omicron has been shown to have more than 40 mutations in Spike protein of SARS-CoV-2 suggesting that the protein is highly amenable to mutations [34]. Additionally, it was also observed that some mutations including 241C>T, 3037C>T, 14408C>T and 23403A>G which originated in February-April 2020 were consistently present till March 2021 suggesting that these mutations might play some critical role in viral pathogenesis.

**Figure 2:**
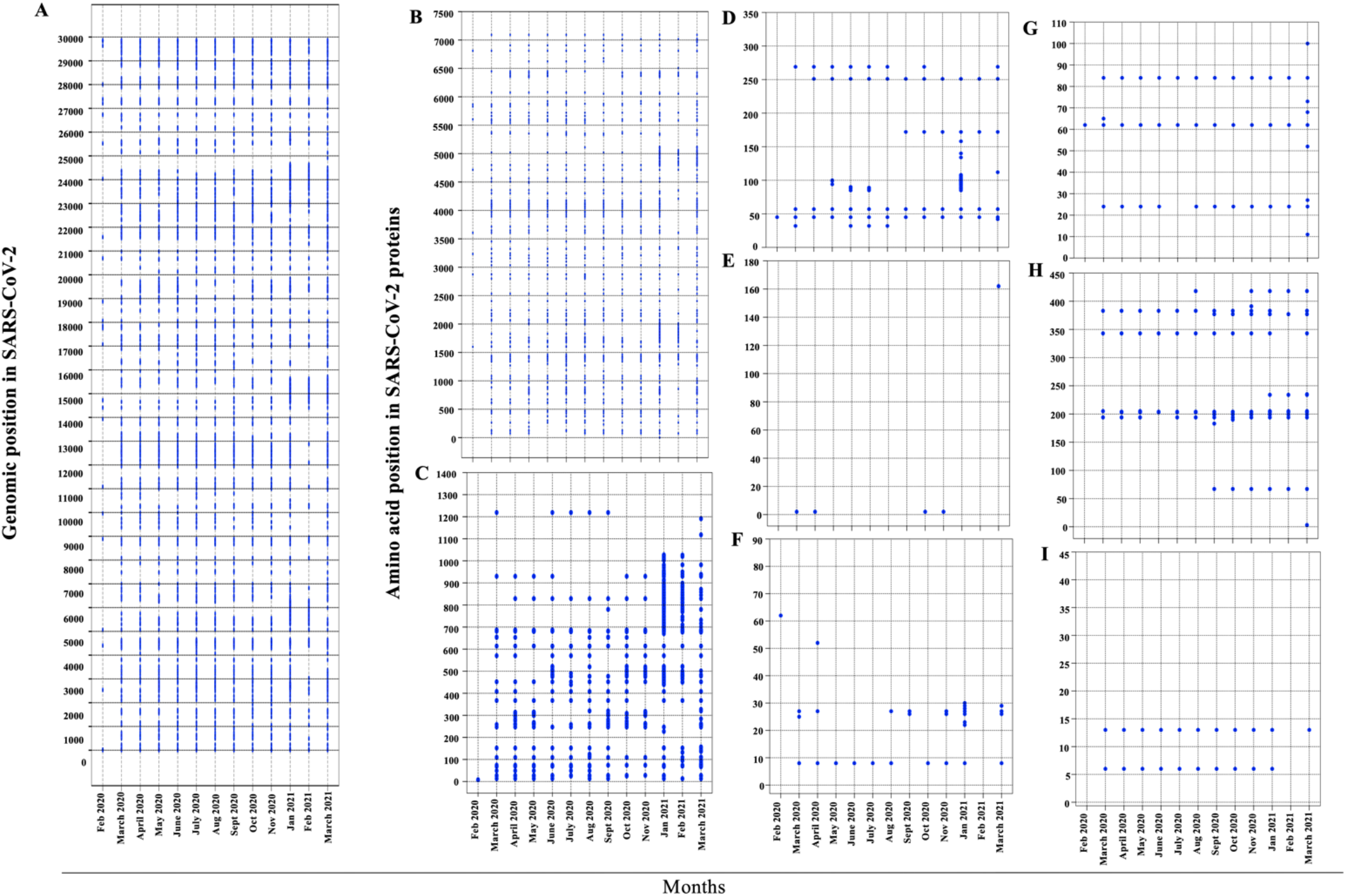
Identification of highly significant mutations occurring in SARS-CoV-2 genomes across various months. (A) Figure showing mutation at nucleotide level among SARS CoV-2 genomes. Figure showing the non-synonymous mutations at amino-acid level in (B) ORF1ab protein (C) S protein (D) ORF3a protein (E) M protein (F) ORF6 protein (G) ORF8 protein (H) N Protein and (I) ORF10 protein of SARS-CoV-2. X-axis represents months whereas Y-axis for represents respective position of SARS-CoV-2.

**Table 2:** Table showing the frequency of unique mutations in various proteins of SARS-CoV-2. The mutation frequency was calculated by dividing the total number of mutations with length of the respective protein.

| SARS-CoV-2 Protein | Length of Protein | Number of Unique Mutation | Frequency |
| --- | --- | --- | --- |
| Orf1ab | 7096 | 610 | 8.59 |
| S | 1273 | 256 | 20.10 |
| Orf3a | 275 | 33 | 12 |
| M | 222 | 2 | 0.90 |
| Orf6 | 61 | 11 | 18.0 |
| Orf8 | 121 | 10 | 8.2 |
| N | 419 | 16 | 3.8 |
| Orf10 | 38 | 2 | 5.1 |

### Correlation among highly significant SARS-CoV-2 mutations

Co-occurrence of several mutations has been shown to modulate the function of the proteins [35]. Therefore, we sought to understand whether there was any association among the mutations that we had identified in this study. For this purpose, we utilized two well-established statistical approaches including Pearson Correlation and Hierarchical Clustering to identify the mutations that were correlated among each other.

### Pearson Correlation

Analysis of SARS-CoV-2 sequences in a time-series manner led us to the identification of several highly significant mutations. In order to identify correlations among mutations in SARS-CoV-2, a binary matrix was created. As discussed earlier, Pearson Correlation was performed on the binary matrix with “1” representing the significant mutation while “0” does not represent the mutations in SARS-CoV-2 genomes. The correlation values range from -0.1 to +0.1 with negative values indicating negative correlation whereas positive values indicating positive correlations. Additionally, the absolute values closer to 1 indicates very strong correlations. Therefore, the results obtained from the Pearson Correlation were then filtered to obtain only those mutations where the absolute value of the correlation coefficient was greater than 0.4. Using this criterion, we obtained 2205 mutation pairs (Figure 3(A)). Though these mutation pairs were highly correlated, however, the frequency of the majority of these mutations was very less. For instance, a correlated mutation pair 21306 and 22995 with absolute correlation >0.4 but occurrence in less than 5% of the genomes might not be of interest. Therefore, to consider only statistically significant pairs of correlated mutations, we filtered the results to retain only those correlated mutations that were present in >30% of the genomes. Using this stringent criterion, we were able to identify 9 mutations (16 mutation pairs) that had an absolute value of correlation > 0.4 and were present in >30% of the genomes (Figure 3(B) and Table 3). It was further observed that six mutation pairs were present in >89% of the genomes suggesting their possible role in viral fitness. Our analysis captured a highly prevalent mutation in Spike protein (D614G) that has nearly replaced the wild-type genome and has been shown to increase the viral infectivity [12]. The identification of D614G mutation using our approach further validated our approach and prompted us to further explore other mutation pairs that were identified.

**Figure 3:**
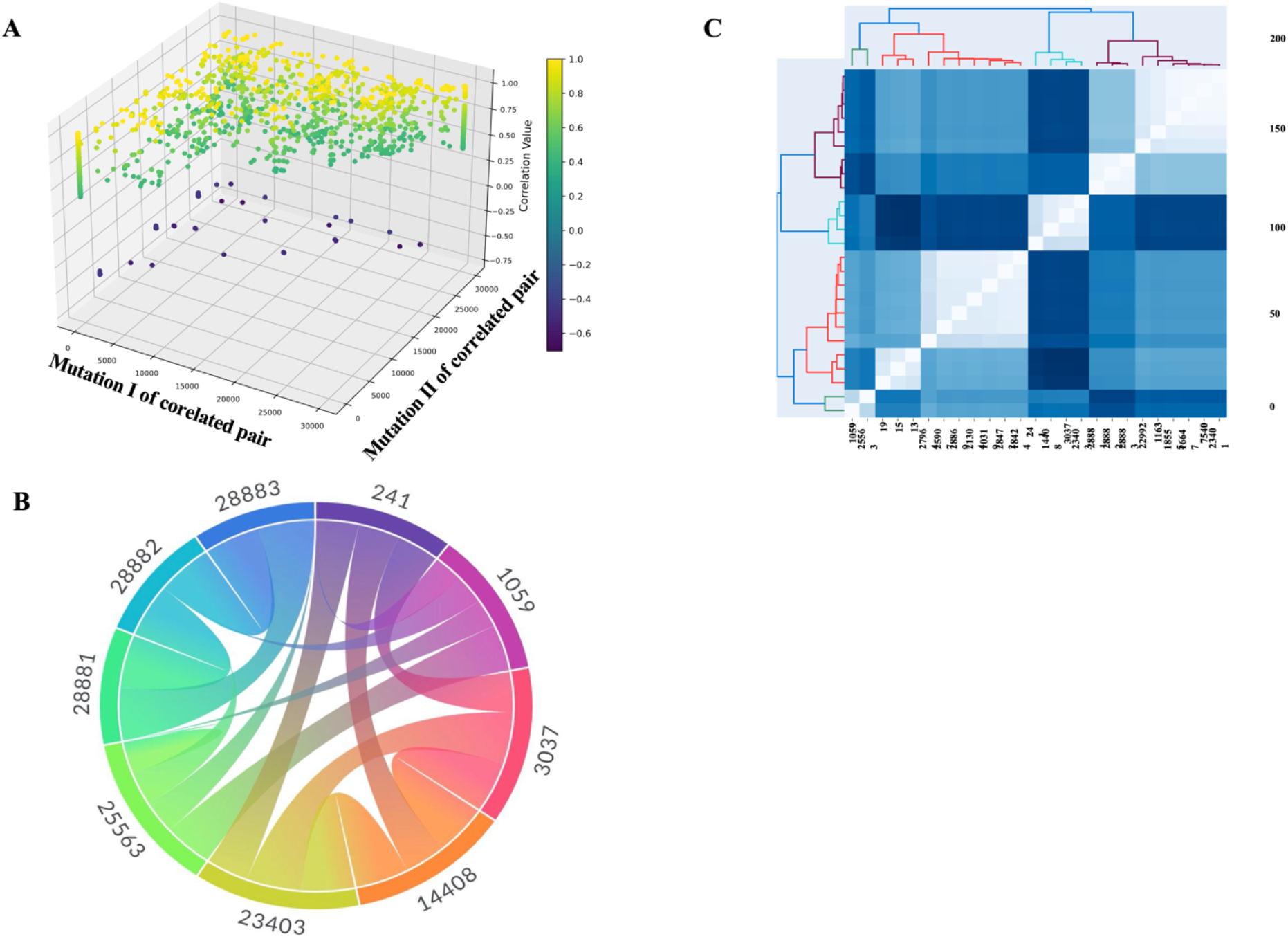
representing correlation and Hierarchical clustering among significant mutations in the genomes of SARS-CoV-2. (A) 3D plot showing the correlation among highly significant mutations having absolute correlation >0.4. X-axis represents mutation I of correlated pair, Y-axis represents mutation II of correlated pair whereas Z-axis represents correlation correlation coefficient. (B) Chord plot representing correlation among significant mutation pairs with absolute correlation value >0.4 and mutation frequency >30% in genomes used in this study (C) Hierarchical clustering of top 25 highly significant mutations having frequency >10%. The color bar represents the distance among the mutations.

**Table 3:** Table showing the correlations among various unique mutations in SARS-CoV-2 genomes. The mutations shown below have correlation value > 0.4 (absolute) and are present in >30% of the genomes.

| Mutation I | Mutation II | Correlation Value | Percentage of genomes that have mutation I | Percentage of genomes that have mutation II |
| --- | --- | --- | --- | --- |
| 241 (5' UTR) | 3037 (F106F NSP3) | 0.760 | 91.054 | 89.889 |
| 241 (5' UTR) | 23403 (D614G Spike) | 0.758 | 91.054 | 90.082 |
| 241 (5' UTR) | 14408 (P323L RdRp) | 0.733 | 91.054 | 89.385 |
| 1059 (T85I NSP2) | 25563 (Q57H ORF3a) | 0.863 | 40.022 | 46.948 |
| 1059 (T85I NSP2) | 28882 (R203R Nucleocapsid) | -0.534 | 40.022 | 30.234 |
| 1059 (T85I NSP2) | 28881 (R203K Nucleocapsid) | -0.535 | 40.022 | 30.381 |
| 1059 (T85I NSP2) | 28883 (G204R Nucleocapsid) | -0.535 | 40.022 | 30.259 |
| 3037 (F106F NSP3) | 23403 (D614G Spike) | 0.976 | 89.889 | 90.082 |
| 3037 (F106F NSP3) | 14408 (P323L RdRp) | 0.943 | 89.889 | 89.385 |
| 14408 (P323L RdRp) | 23403 (D614G Spike) | 0.943 | 89.385 | 90.082 |
| 25563 (Q57H ORF3a) | 28882 (R203R Nucleocapsid) | -0.613 | 46.948 | 30.234 |
| 25563 (Q57H ORF3a) | 28881 (R203K Nucleocapsid) | -0.614 | 46.948 | 30.381 |
| 25563 (Q57H ORF3a) | 28883 (G204R Nucleocapsid) | -0.614 | 46.948 | 30.234 |
| 28881 (R203K Nucleocapsid) | 28882 (R203R Nucleocapsid) | 0.995 | 30.381 | 30.259 |
| 28881 (R203K Nucleocapsid) | 28883 (G204R Nucleocapsid) | 0.994 | 30.381 | 30.259 |
| 28882 (R203R Nucleocapsid) | 28883 (G204R Nucleocapsid) | 0.997 | 30.234 | 30.259 |

### Hierarchical Clustering

In order to garner confidence in our approach, we used another widely used statistical approach to identify the clusters among highly significant mutations in SARS-CoV-2. Since Hierarchical Clustering is a computationally intensive process, we analysed only the top 25 mutations that were highly significant and were present in >10% of the genomes used in this study (Table 4). We obtained a binary matrix for the top 25 highly significant mutations. Similar to the results obtained using Pearson Correlation, Hierarchical Clustering analysis led to clustering of the similar mutations. Here we analyzed only those mutation clusters that were present in greater than 30% of genomes. (Figure 3(C)). It can be observed that 241C>T, 14408C>T, 3037C>T and 23403A>G forms a cluster and are most common concurrent mutations followed by 28881G>A, 28882G>A and 28883G>C. Since both the statistical tools provided similar results, we then focussed our attention to these mutations for their in-depth understanding.

**Table 4:** showing the mutations and their frequency. NC refers to the total number of sequences with a specific mutation.

| Mutation | Gene | NC |
| --- | --- | --- |
| 241 | 5'UTR | 50771 |
| 23403 (D614G) | S | 50229 |
| 3037 (F924F) | ORF1AB | 50121 |
| 14408 (P4715L) | ORF1AB | 49840 |
| 25563 (Q57H) | ORF3A | 26178 |
| 1059 (T265I) | ORF1AB | 22316 |
| 28881 (R203K) | N | 16940 |
| 28883 (G204R) | N | 16872 |
| 28882 (R203R) | N | 16858 |
| 27964 (S24L) | ORF1AB | 10985 |
| 1163 (I300F) | ORF1AB | 9098 |
| 10319 (L3352F) | ORF1AB | 8882 |
| 18555 (D6097D) | ORF1AB | 8772 |
| 28869 (P199L) | N | 8770 |
| 16647 (T5641T) | Helicase | 8755 |
| 23401 (Q613H) | S | 8746 |
| 7540 (T2425T) | ORF1AB | 8729 |
| 18424 (N6054D) | ORF1AB | 8758 |
| 28472 (P67T) | N | 8572 |
| 21304 (R7014S) | ORF1AB | 8320 |
| 25907 (G172D) | ORF3A | 8236 |
| 22992 (S477N) | S | 8146 |
| 19 | 5'UTR | 6637 |
| 15 | 5'UTR | 5662 |
| 13 | 5'UTR | 5069 |

### Frequency and global distribution of highly, correlated and frequent significant mutations

Once the correlated pair of highly significant mutations that have absolute correlation coefficient >0.4 and present in >30% of genomes were identified, we then investigated their global distribution among the circulating SARS-CoV-2 genomes. It can be observed that C241T mutation in the 5’UTR region completely replaced the wild-type genomes as early as June-July 2020 (Figure 4). Similar trends were observed with the mutations C3037T, C14408T, and A23403G suggesting their critical role in viral pathogenesis. However, some of the mutations showed a mosaic pattern of global distribution with increase in time.

**Figure 4:**
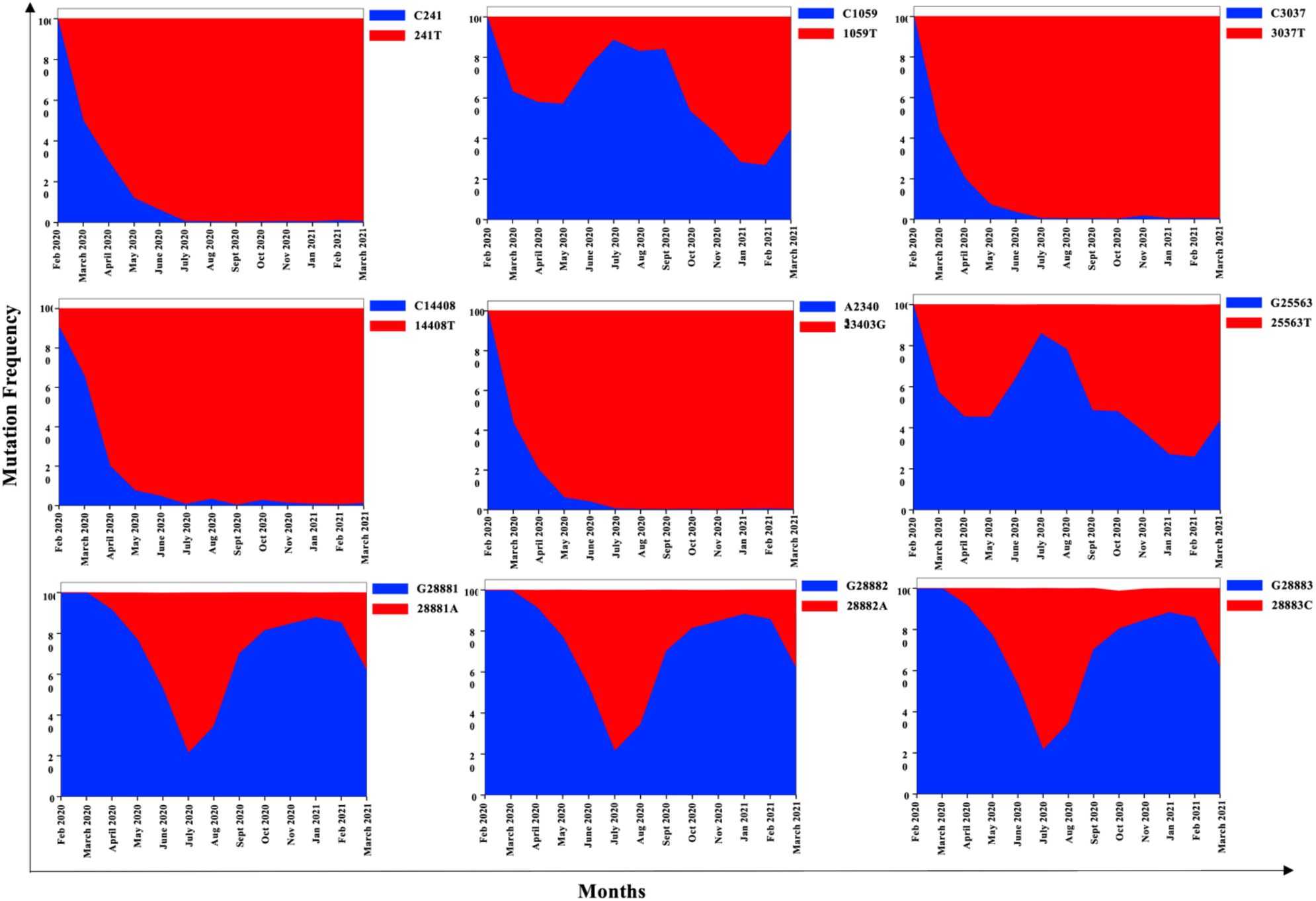
showing the running monthly counts of the mutations in SARS-CoV-2 genomes used in this study. The highly significant mutations that are correlated with each other shown here are present in >30% of the genomes and have absolute correlation coefficient >0.4 (A) 241C>T (B) 1059C>T (C) 3037C>T (D) 14408C>T (E) 23403A>G (F) 25563G>T (G) 28881G>A (H) 28882G>A (I) 28883G>C.

### Mutation in the 5’ UTR

The untranslated region of the viral genome plays a vital role in viral replication. The region has been shown to form various secondary structures to allow binding of cellular and viral proteins thereby regulating the translation of viral proteins [36,37]. Therefore, any mutation in these highly conserved regions has the potential to regulate viral replication. Statistical approaches revealed that 241C>T mutation was highly correlated with three different mutations 3037C>T (F106F), 14408C>T (P323L) and 23403A>G (D614G) in SARS-CoV-2 Nsp2, RdRp and Spike proteins respectively. Remarkably, it can be observed that the correlation of mutation 241C>T with all the other mutations mentioned above was >0.75 pointing towards a very strong correlation. Additionally, these mutations were found in > 89% of the genomes further pointing towards the critical role of these mutations in viral evolution. These observations were further supported by Hierarchical Clustering where these mutations were clustered together. Our results are in agreement with published studies which have shown similar correlations among these mutations [14]. However, these studies were conducted on the genomes from various countries including Israel, USA, and Bangladesh whereas our analysis is obtained from the SARS-CoV-2 genomes obtained globally. The correlation of 241C>T mutation with highly occurring mutations in the SARS-CoV-2 genomes point towards its role in viral pathogenesis and fitness.

### Mutation in the Nsp2 protein

The Nsp2 protein of SARS-CoV-2 was recently shown to be associated with host proteins involved in vesicle trafficking. It was also proposed that targeting viral protein N, Nsp2 and Nsp8 interactions with host translational machinery might have therapeutic effects [38]. Therefore, understanding the dynamics of Nsp2 protein becomes essential. Our analysis revealed that mutation 1059C>T (T85I) in viral Nsp2 protein was both positively and negatively correlated with other mutations in SARS-CoV-2. As described in Table 3, 1059C>T (T85I) mutation in Nsp2 protein was positively correlated with 25563G>T (Q57H) mutation in ORF3a. The correlation coefficient is 0.863 and presence of these mutations in >40% of the genomes suggests that the co-occurrence of these mutations might play a role in viral evolution. These observations are in agreement with earlier studies where co-occurrence of 1059C>T (T85I) with 25563G>T (Q57H) was observed in nearly 70% of COVID cases across the USA [15].

Additionally, we observe that 1059C>T (T85I) mutation in Nsp2 protein was negatively associated with three mutations 28881G>A (R203K), 28882G>A(R203R) and 28883G>C (G204R) in the N protein. Though the co-occurrence of these mutations is established in other study also [14], however, in this study, we showed that these mutations are negatively correlated from the SARS-CoV-2 genomic sequences across the world.

Since T85I mutation was widespread among Nsp2 protein, we sought to investigate the role of this mutation on the protein function. The full-length 3.2 Å crystal structure of Nsp2 (PDB: 7SMW) was solved by combining cryo-EM and recently developed AI tool AlphaFold2 [39]. In the structure, there was a highly conserved zinc binding site observed that indicates the role of Nsp2 in RNA binding. Figure 5(A) represents the structure of Nsp2. In Nsp2, T85I mutation in which polar threonine residue is substituted by hydrophobic isoleucine at position 85 at β-hairpin loop formed by two β-strands (1. 78-82 a.a. & 2. 98-104 a.a.). PredictSNP revealed that this mutation is deleterious in nature with around a 70% confidence score. The ENCoM based ΔΔS_vib_ value suggests that mutation brings some extent of flexibility in Nsp2 protein. It can be seen from Figure 6(A) that two helices (1: 19-28 a.a. & 2: 35-45 a.a.) at the N-terminus gain slight flexibility (red colour). Among six predictors, four predicted ΔΔG negative that implies destabilization in Nsp2 (Table 5). Our results on Nsp2 stability and flexibility are in accordance with already published reports [15]. In WT and MT, two identical residues (Phe83 & Asn87) interact with WT and MT residues. In both the structures, van der Waals clashes were observed between sidechain oxygen of Thr85 in WT and aromatic carbons of Phe83 and, sidechain methyl group carbon atom of Ile85 in MT and aromatic carbons of Phe83. In WT, Thr85 amide group oxygen and nitrogen interact with surrounding amide group atoms of Phe83 and Asn87 through hydrophobic, vdW and polar interactions. However, in MT, similar interactions were noted but, there was a polar-vdW clash observed between Asn87 and Ile85. This might be the cause of the predicted instability of T85I mutation in Nsp2 protein. WT and MT interactions are illustrated in Figure 6(D).

**Figure 5:**
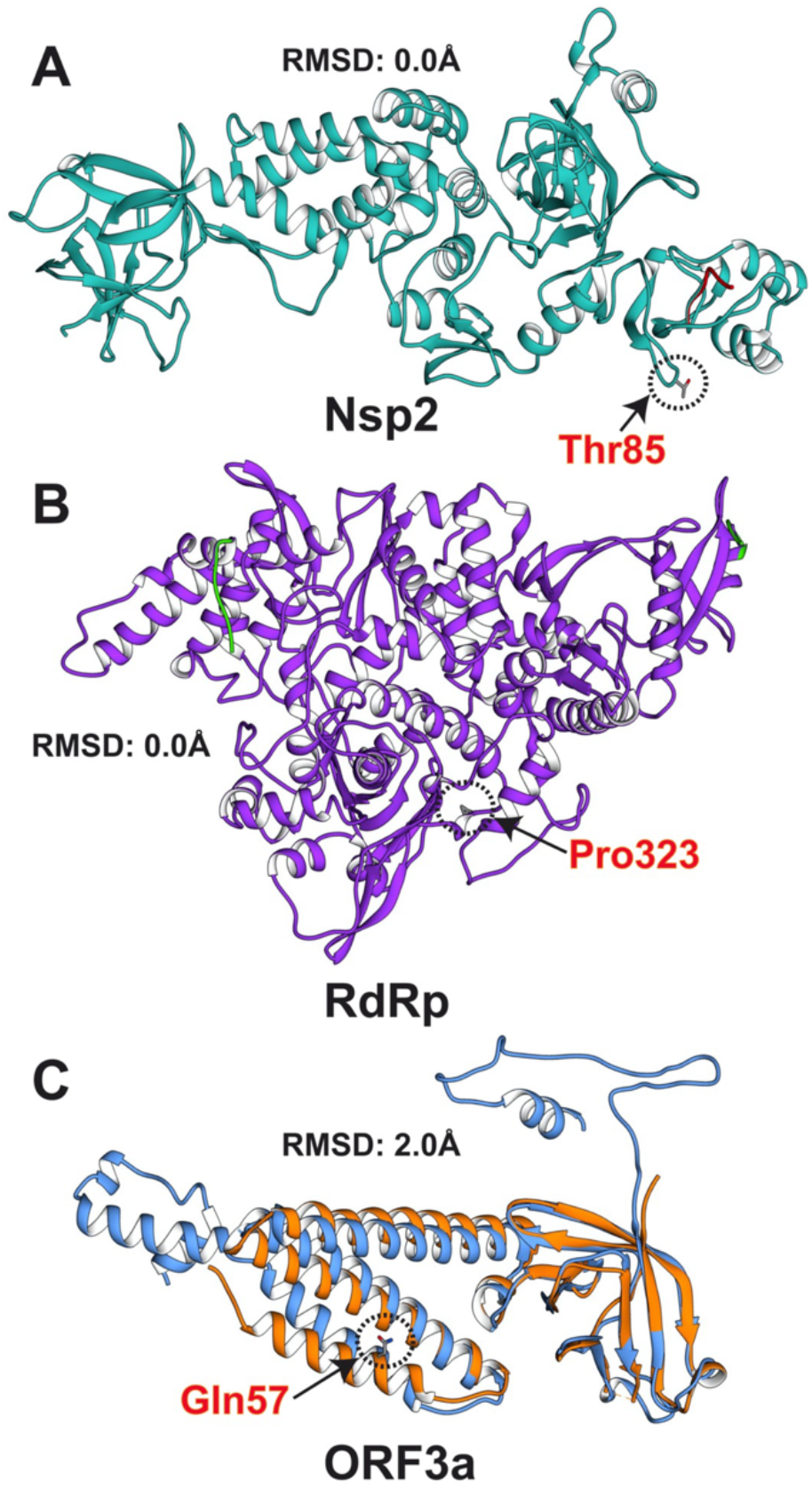
Structure alignments of crystal and modelled proteins. (A) Nsp2. Crystal and modelled structures are cyan and red in color respectively. (B) RdRp. Crystal and modelled structures are purple and red in color respectively. (C) ORF3a. Crystal and modelled structures are orange and blue in color respectively. Mutation positions are indicated in dashed circle and mutant residues are represented in stick form.

**Figure 6:**
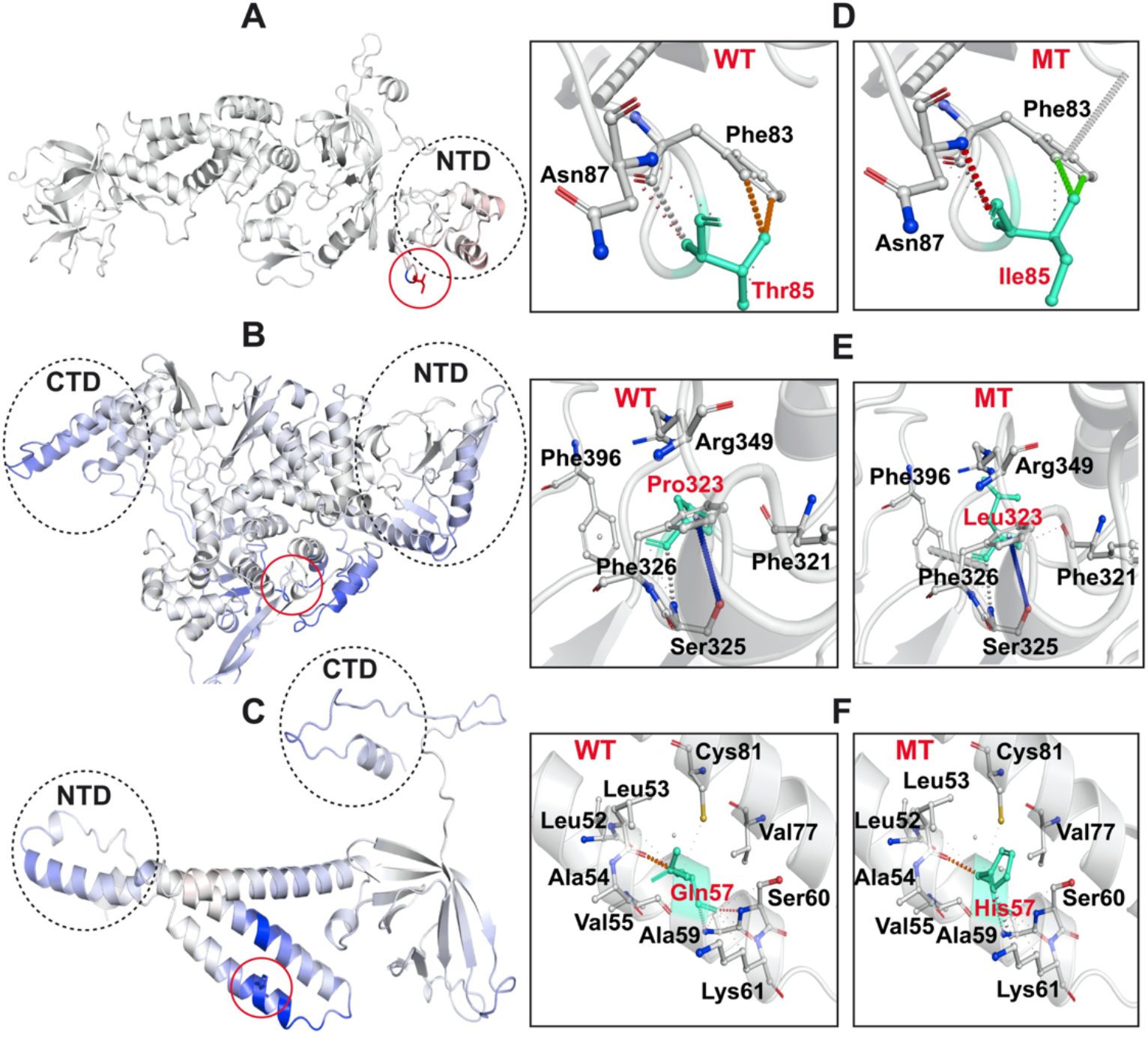
Visual representation of MT protein dynamicity and intramolecular interactions of WT and MT residues with proximal amino acids. Mutations are shown in sticks representation and are inside the red circle. Red and blue colours indicate the flexibility and rigidity respectively. WT and MT residues are shown in cyan colour. Mutant residues are red in color while interacting residues are in black color. Interactions are illustrated in different colours and for the further interpretation of interactions, readers are requested to visit the Web version of Arpeggio web server. (A-C) Visual representation of Nsp2, RdRp, and ORF3a respectively. (D-F) Intramolecular interactions of WT and MT residues in Nsp2, RdRp, and ORF3a respectively.

**Table 5:**
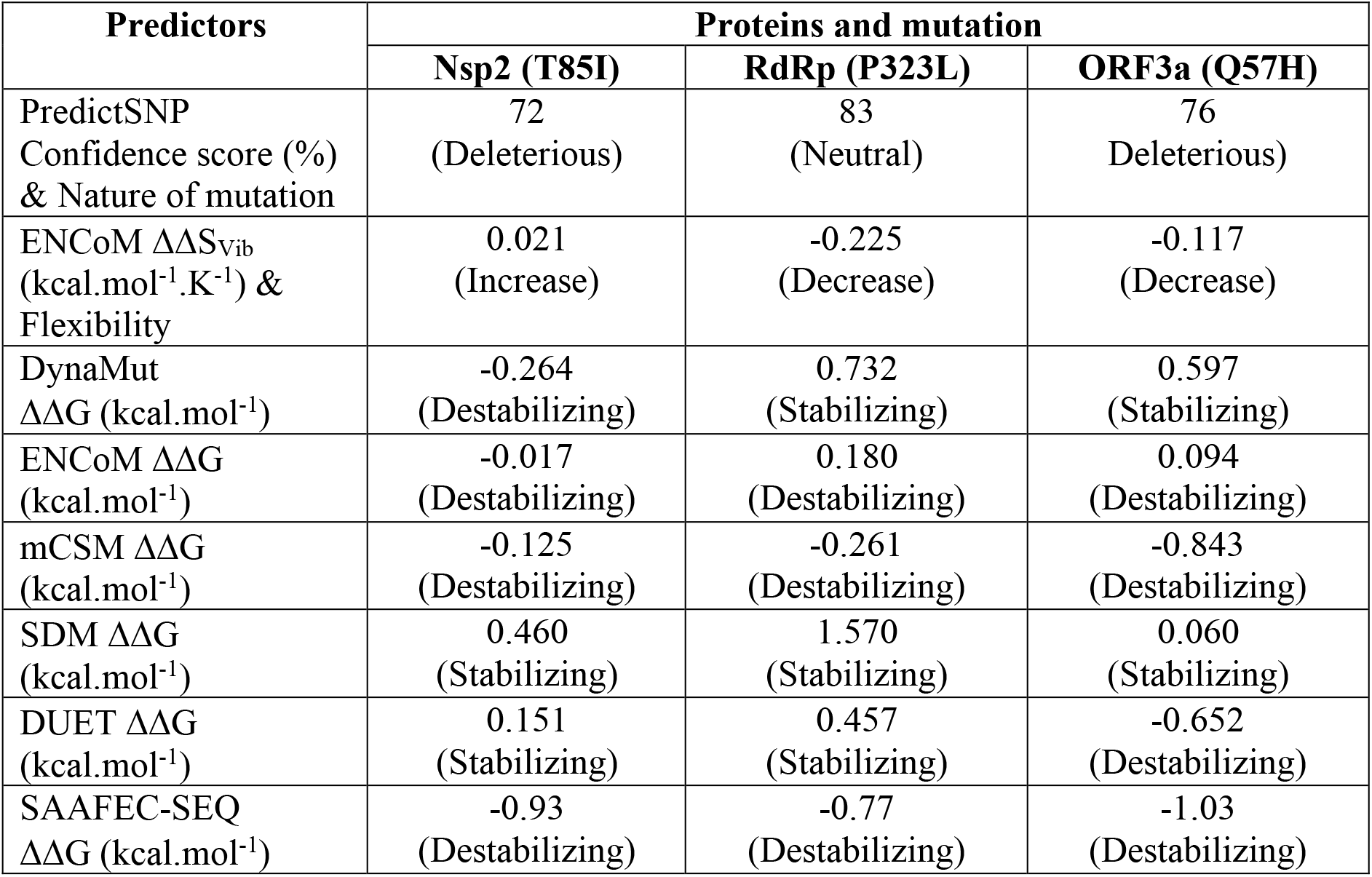
Predicted results for the effects of mutation on functionality, stability and flexibility of proteins using PredictSNP, DynaMut and SAAFEC-SEQ web servers.

| Predictors | Proteins and mutation |  |  |
| --- | --- | --- | --- |
|  | Nsp2 (T85I) | RdRp (P323L) | ORF3a (Q57H) |
| PredictSNP<br>Confidence score (%)<br>& Nature of mutation | 72<br>(Deleterious) | 83<br>(Neutral) | 76<br>(Deleterious) |
| ENCoM $\Delta\Delta S_{vib}$<br>(kcal.mol<sup>-1</sup>.K<sup>-1</sup>) &<br>Flexibility | 0.021<br>(Increase) | -0.225<br>(Decrease) | -0.117<br>(Decrease) |
| DynaMut<br>$\Delta\Delta G$ (kcal.mol<sup>-1</sup>) | -0.264<br>(Destabilizing) | 0.732<br>(Stabilizing) | 0.597<br>(Stabilizing) |
| ENCoM $\Delta\Delta G$<br>(kcal.mol<sup>-1</sup>) | -0.017<br>(Destabilizing) | 0.180<br>(Destabilizing) | 0.094<br>(Destabilizing) |
| mCSM $\Delta\Delta G$<br>(kcal.mol<sup>-1</sup>) | -0.125<br>(Destabilizing) | -0.261<br>(Destabilizing) | -0.843<br>(Destabilizing) |
| SDM $\Delta\Delta G$<br>(kcal.mol<sup>-1</sup>) | 0.460<br>(Stabilizing) | 1.570<br>(Stabilizing) | 0.060<br>(Stabilizing) |
| DUET $\Delta\Delta G$<br>(kcal.mol<sup>-1</sup>) | 0.151<br>(Stabilizing) | 0.457<br>(Stabilizing) | -0.652<br>(Destabilizing) |
| SAAFEC-SEQ<br>$\Delta\Delta G$ (kcal.mol<sup>-1</sup>) | -0.93<br>(Destabilizing) | -0.77<br>(Destabilizing) | -1.03<br>(Destabilizing) |

### Mutations in the Nsp3 protein

The Nsp3 protein in coronaviruses has been shown to antagonize the innate immune responses [40]. The mutations in the Nsp3 macrodomain region lead to enhanced type I IFN responses and reduced viral replication [41]. Understanding the dynamics of mutations in Nsp3 might provide clues on SARS-CoV-2 mediated evasion of type I IFN signalling. We identified a synonymous mutation 3037C>T (F106F) in SARS-CoV-2 Nsp3 protein. Though silent in nature, this mutation was shown to disrupt the mir-197-5p target sequence [42]. Mir-197-5p was shown to be associated with some other viruses also [43–45] thereby pointing towards its role in viral biology. This mutation was highly correlated with various mutations spread across the genome of SARS-CoV-2. It was positively correlated with 241C>T in the 5’UTR region, 23403A>G (D614G) in S protein and 14408C>T (P323L) mutation in RNA dependent RNA polymerase (RdRp). These mutation correlation pairs have been shown to be critical in evolution to the more infectious form of SARS-CoV-2. The 14408 mutation was shown to increase the mutation rate among SARS-CoV-2 whereas D614G has been shown to contribute towards the infectivity of the virus [16]. The presence of all these mutations in >90% of the genomes further points towards their critical role in driving viral evolution.

### Mutation in the RdRp protein

The RdRp (Nsp12) protein of the SARS-CoV-2 is the RNA-dependent RNA-polymerase which is important for viral replication and transcription. This protein is also believed to be the most prominent target for potential antiviral drugs [46]. Therefore, understanding the mutations in this protein becomes critical for RdRp based drug designs. We identified a highly significant mutation 14408C>T (P323L) in this protein which was present in >89% of genomes suggesting that the mutation was now a part of the circulating genomes. Apart from its widespread presence, this mutation was strongly associated with some other mutations including 241C>T in 5’UTR, 3037C>T in Nsp3, and 23403A>G in S with extremely high absolute correlation values 0.73, 0.94, and 0.94 respectively. Interestingly, P323L and G614G mutations were reported in severe COVID-19 patients as compared to the mild ones suggesting their possible role in viral pathology [47]. Owing to the widespread presence 14408C>T (P323L) mutation in RdRp, we sought to study its putative effect of the stability of wt RdRp. The 2.83 Å crystal structure of RdRp in complex with Nsp7, Nsp8, Nsp9, and Helicase was determined using cryo-EM [48]. RdRp structure has the RdRp domain (367-920 a.a.) [49] and N-terminus (60-249 a.a.) which adopts Nidovirus RdRp-associated nucleotidyltransferase (NiRAN) structural scaffold [50]. Another region (4-118 a.a.) is composed of two helices and five β-strands that are antiparallel. Additionally, the short β-strand (215-218 a.a.) was observed in RdRp which is highly ordered in SARS-CoV-2 as compared to SARS-CoV. This β-strand has contact with other β-strand residues (96-100 a.a.) consequently, it increases the conformational stability of RdRp in SARS-CoV-2 [49]. The structure of RdRp is given in Figure 5(B).

P323L mutation is present on RdRp interface domain and, especially in the loop region which connects the interface domain’s three helices to the same domain’s three β-strands. Earlier study suggests that this mutation enhances the processivity of RdRp protein [51]. It is predicted functionally neutral with a notable confidence score (83%) to the protein. In this mutation, a conformationally rigid proline ring is swapped by a flexible side-chain containing leucine residue. Though the nature of WT and MT residues is hydrophobic, their conformational flexibility must be the deciding factor for the protein stability and flexibility. Nonetheless, this mutation significantly rigidifies the RdRp (Figure 6(B)) and ΔΔS_vib_ also observed least (Table 5). Results show that this mutation has a strong communication network in RdRp and impacts various helices and β-sheets. P323L mutation is located in a loop formed by the β-strand (328-335 a.a.) and helix (304-320 a.a.) thus, these two secondary structures gain rigidity. However, a helix nearby mutation gained greater rigidness as compared to other regions of the RdRp. The helices at the N- and C-terminal domains also gained rigidness due to this mutation. All-atom simulation data also suggested that P323L mutation reduces the RdRp motions thus, this is in line with our results [52]. The ΔΔG value prediction showed that three predictors predicted stabilization and the remaining three predicted destabilization but the ΔΔG values of stabilization are considerably higher in comparison with destabilization values. Mohammad et al. have performed 200 ns all-atom MD simulation and by calculating free energy (ΔG) of WT and MT RdRp they confirmed that P323L mutation increases the stability of MT RdRp [52]. Hence, this mutation stabilizes the RdRp structure.

The RdRp WT and MT interactions analysis revealed that there are a greater number of interactions observed in MT as compared to WT. The WT and MT residues are surrounded by Phe321, Ser325, Phe326, Arg349 and Phe396 residues. In WT, only single polar interaction between Ser325 and Pro323 residues whilst in MT, two additional hydrogen bonds were observed through Ser325 and Phe326 and polar interaction through Phe349. Thus, it can be considered that higher stability in MT comes from these interactions. Figure 6(E) shows the interactions in WT and MT RdRp proteins.

### Mutation in the ORF3a protein

The largest accessory protein ORF3a plays a key role in the viral infection cycle. Moreover, this protein is essential for viral replication, and mutations in this protein are associated with higher mortality rates [53]. We identified a highly significant mutation 25563G>T (Q57H) in SARS-CoV-2 ORF3a protein. Interestingly, this mutation has been shown to be associated with decreased death and increased cases of COVID-19 [54]. We further show that 25563G>T (Q57H) mutation is positively associated with 1059C>T mutation in Nsp2 protein whereas it is negatively associated with 28881G>A, 28882G>A, 28883G>C mutations in N protein. Our observations are in agreement with previous studies which have identified similar associations within genomes of SARS-CoV-2 isolated from Israel [14].

The ORF3a functional domains are vital for SARS-CoV-2 infectivity, virulence, ion channel synthesis and release of virus [55]. Previous study showed that ORF3a in SARS-CoV-2 has a weaker potential for pro-apoptotic activity as compared to SARS-CoA ORF3a. The difference between pro-apoptotic activity might be linked to the infectivity of viruses [56]. Furthermore, another study has confirmed that ORF3a binds to the Homotypic fusion and protein sorting (HOPS) and prevents the autolysosome formation [57]. ORF3a is also considered potential vaccine and drug targets [58,59]. The experimental structure of ORF3a (PDB: 6XDC) was determined using cryo-EM at 2.1 Å resolution. ORF3a has three main regions, N-terminal (1-39 a.a.), cytoplasmic loop (175-180 a.a.) and C-terminal (239-275 a.a.) [60]. Figure 5(C) shows the structure of ORF3a. Finally, the Q57H mutation in ORF3a was studied. In this mutation, glutamine polar residue is exchanged by polar and basic histidine residue. This mutation is situated in the helix region of ORF3a. This mutation has deleterious nature with a 76% confidence score (Table 5) thus, it might alter the functions of ORF3a protein. This mutation was reported predominantly occurring and deleterious in the previous studies [55,61].

The vibrational entropy energy (ΔΔS_vib_) value was noted negative hence, ORF3a gains rigidness and becomes less flexible due to the insertion of this mutation. Similar to RdRp, mutant residue in ORF3a has also wide communication dynamics. This single mutation at helix increases the rigidity of the whole ORF3a protein (Figure 6(C)). Our finding supports previous study on Q57H mutation and its rigidness [15]. This mutation was predicted destabilizing by the majority of predictors based on ΔΔG values (Table 5). WT and MT residues are in close proximity to Leu52, Leu53, Ala54, Val55, Ala59, Ser60, Lys61, Val77 and Cys81. In WT, two hydrogen bonds are observed by Ser60 and Lys61 amide bond amino groups with amide carbonyl group of WT residue. These identical hydrogen bonds are also present in MT structure. Other types of interactions such as polar and hydrophobic were observed in WT and MT ORF3a. But there are two new clashes seen in MT structure between the histidine ring and Lys61 and mutant amide group and Leu53 residue. Thus, the overall number of clashes has increased in MT and this might be the leading factor of destabilization of ORF3a protein under the influence of Q57H mutation. This mutation was predicted to be having significant potential to alter the ORF3a conformations and leads to disruption of intramolecular hydrogen bonds in ORF3a [54]. Thus, our finding matches that Q57H mutation causes destabilization of ORF3a with this study. WT and MT interactions are illustrated in Figure 6(F).

### Mutation in the Spike protein

Spike protein is a homotrimer protein that is studded on the surface of SARS-CoV-2 thereby giving it a crown like shape. Spike protein of SARS-CoV-2 is cleaved in the infected cells and consists of two subunits that are covalently attached to each other. One of the subunits i.e S1 binds to the ACE2 receptor on the target cells where the other unit S2 helps in membrane fusion a transmembrane protein that binds with the receptors on host cells to enable the virus to enter inside the cells [62,63]. D614G mutation in the spike proteins has been shown to increase the infectivity of the virus. In our previous study [20], we also characterized that the effect of D614G mutation on protein activity and suggested that the mutation led to decreased protein stability but enhanced protein movement. In the present study, we obtained a correlation of D614G mutation in spike protein with mutations in 5’UTR (241C>T), Nsp3 protein (3037C>T) and RdRp (14408C>T). The presence of these co-mutation pairs in >96% points towards the critical role that these mutations play in viral fitness.

### Mutation in the Nucleocapsid protein

Nucleocapsid protein is one of the most conserved proteins among SARS coronaviruses. This protein is known to interact with viral RNA as well as M protein to aid virion assembly. This protein is also shown to play a role in regulating host immune responses [64] and cellular apoptosis [65]. SARS-CoV-2 Nucleocapsid protein acts as a viral RNAi suppressor where it has been shown to antagonize cellular RNAi pathways [66]. Thus, understanding the role of mutations in modulating the function of this protein becomes important. Our statistical analysis and time series data analysis identified positive correlation of 28881G>A, 28882G>A and 28883G >C with mutations in Nsp2 (1059C>T) and ORF3a (25563G>T) respectively. These three mutations are correlated with each other. Among the three mutations in the Nucleocapsid protein, mutations at 28881 and 28883 are missense mutations whereas 28882 is a synonymous mutation. Since the mutations in nucleocapsid protein are known to increase the infectivity and virulence of the virus [67], therefore, the correlation of these mutations with other mutations warrants further study. In our recent report [20] we used computational tools to characterize the mutation in N protein and have reported that mutation at 28881 (R203K) stabilizes the parent protein whereas at 28883 (G204R) destabilizes the N protein.

### Impacts of mutations on protein dynamics

The protein structure and functions are significantly altered by the insertion of single point mutations [68–70]. Investigating the structural or functional impacts of point mutations in all protein is a mammoth task, thus, computational algorithms and tools like NMA models [71], Gaussian network models (GNM) [72], Anisotropic network models (ANM) [73], Elastic network models (ENM) [74], Discrete molecular dynamics (DMD) [75], All-atom molecular dynamics (AAMD) simulation [76] and protein evolutionary data are being used currently. Therefore, we employed numerous computational tools to probe the effects of mutations on protein structures. The predicted results of the mutations are given in Table 5.

### Linear mutual information (LMI) analysis

The normalized LMI gives insight into protein residue network and dynamic correlation. Figure 7 illustrates the nLMI correlation and correlation difference plots of Nsp2, RdRp and ORF3a proteins and their mutants. It can be seen from Figure 7(A-C), that Nsp2 and ORF3a protein residues are strongly correlated (>0.625) as compared to RdRp where correlation among residues is considerably less (<0.500). However, to understand the impacts of every single mutation in each protein, we obtained correlation differences between the WT and MT structures of all three proteins. The correlation difference plots of proteins are shown in Figure 7(D-F). The yellow regions indicate no correlation of slight correlation (0.00-0.25) whereas cyan regions are designated for slightly negative anticorrelation between the residues. In RdRp and ORF3a MTs structures, residues have significantly less correlation. However, Nsp2 MT structure’s residues are slightly anticorrelated. Thus, T85I mutation in Nsp2 causes a slight disruption in the structure that can be considered destabilization of Nsp2 MT structure. But, P323L in RdRp and Q57H in ORF3a do not bring notable destabilization to their mutant structures.

**Figure 7:**
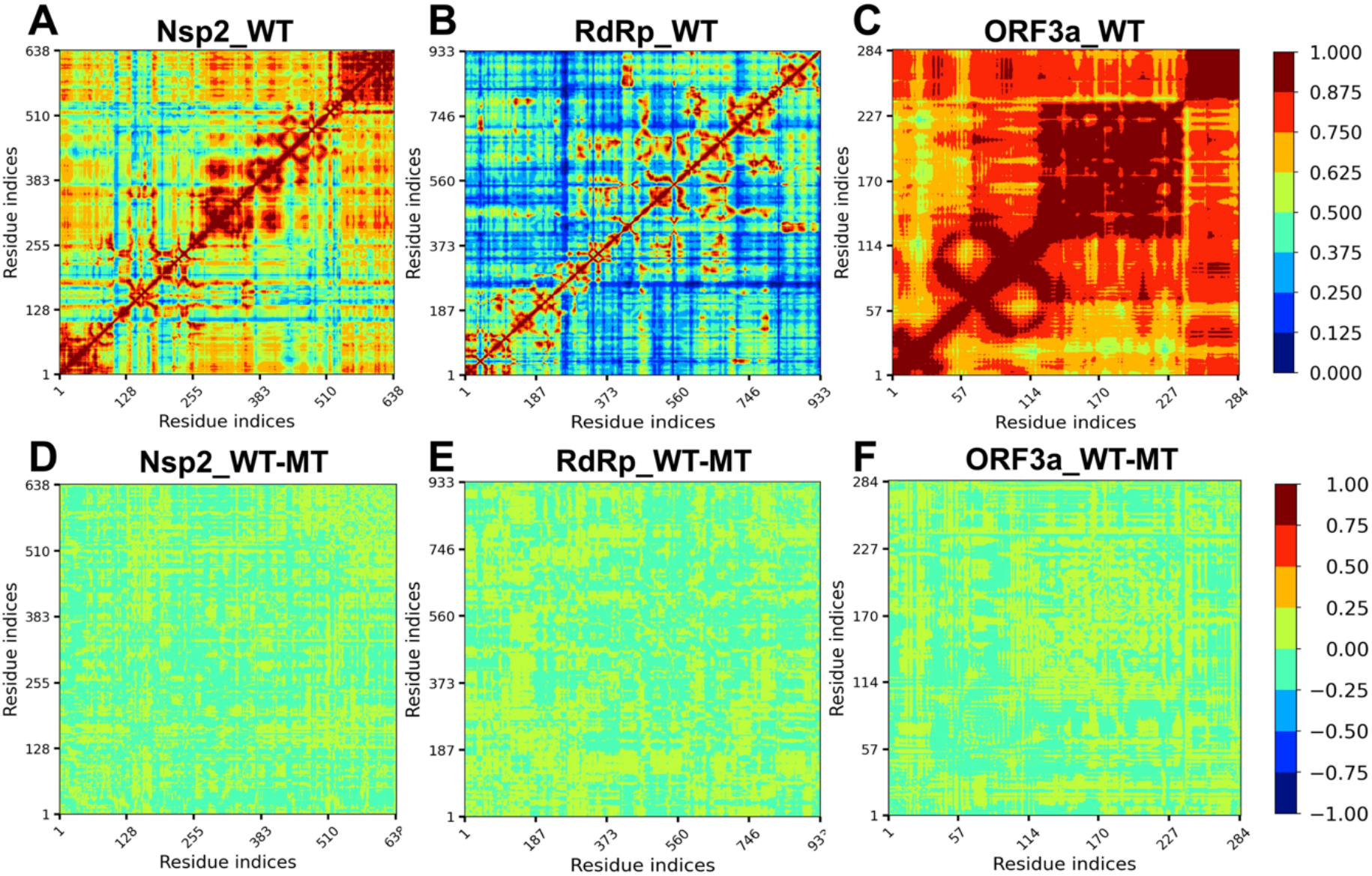
Normalized linear mutual information (nLMI) correlation plots. (A) Nsp2_WT (B) RdRp_WT (C) ORF3a_WT (D) Correlation difference plot of Nsp2 WT & MT (E) Correlation difference plot of RdRp WT & MT (F) Correlation difference plot of ORF3a WT & MT. Degree of correlation corresponds to the color bar.

## Conclusion

Since the onset of the SARS-CoV-2 pandemic in December 2019, the virus has significantly mutated. The mutations in the virus have led to the emergence of mutants that have the capacity to dodge vaccine and antiviral therapies. It now seems that the virus is evolving to be more infectious and less virulent. Therefore, understanding the dynamics of mutations in the viral genome is of utmost importance. In this regard, we performed time-series analysis of viral genomes to understand the origin and frequency of significant mutations that are significant in the SARS-CoV-2 genome. Additionally, we used Pearson Correlation and Hierarchical Clustering to identify correlations among highly significant mutations that are correlated. We identified sixteen mutation pairs that had absolute correlation value >0.4 and were present in >30% of the genomes analysed in this study. We identified a strong correlation coefficient (>0.73) of mutation 241 in 5’UTR (>0.73) of mutation at 241 position in the 5’ UTR with 3037 (F106F) in Nsp3, 23403 (D614G) in Spike, and 14408 (P323L) in RdRp respectively. These mutations were found in >89% of the genomes that were studied.

To investigate the impacts of T85I, P323L, and Q57H mutations in Nsp2, RdRp, and ORF3a respectively on their structural stability and flexibility, we employed structure and sequence-based tools. The free energy change (ΔΔG) and vibrational entropy energy (ΔΔS_vib_) terms were used to evaluate the stability and flexibility of proteins respectively. Results showed that out of three, two mutations (T85I in Nsp2 & Q57H in ORF3a) were predicted to destabilize while P323L was stabilizing. Also, consensus predictors were used to predict the impacts of mutation on protein functions. It was noted that T85I in Nsp2 and Q57H in ORF3a were found to have deleterious nature which implies that they alter the protein functions. However, P323L in RdRp was predicted neutral which suggests that this mutation does not have any impacts on protein function. The graph theory-based nLMI correlation was also obtained for the WT and MT structures of three proteins to understand the residue communication in proteins. The Nsp2 and ORF3a residues have greater correlation in comparison with RdRp. Correlation difference analysis suggests that T85I in Nsp2 is destabilizing whereas P323L in RdRp and Q57H in ORF3a are slightly stabilizing. The fact that some of the mutations have destabilizing effects but have very high frequency suggests that these might be playing some role in viral fitness. Further experimental evidence is required to study the effect of these co-mutations on viral transmission and pathogenesis.

## Acknowledgements

NP is thankful to UGC for PhD fellowship. VS received research grant from UGC, Govt. of India. SBR is thankful to his Chemistry Department for providing computational and infrastructure facilities.

## Author contribution

NP, SS, SBR, AJ, GS performed the experiments and analyzed the data. BK contributed to the statistical experiments. VS, PA, BK, RPB, SBR, KRS designed and supervised the study. NP, SBR, SS, and VS wrote the first draft. VS, PA, BK, SBR, RPB and KRS edited and finalized the draft.

